# Membrane Geometry Reveals Strengths and Limitations of AlphaFold-Predicted Transmembrane Protein Complexes

**DOI:** 10.64898/2026.11.09.765245

**Authors:** Dominik Sziklai, Zoltán Wiener, Rita Padanyi, Tamás Hegedűs

## Abstract

High-confidence protein complex predictions do not necessarily represent physically or biologically plausible assemblies. This limitation is particularly relevant for transmembrane (TM) proteins, whose interactions are constrained by the lipid bilayer. Here, we systematically evaluated membrane compatibility in the recently expanded AlphaFold Protein Structure Database, analysing 189,469 high-confidence AlphaFold-Multimer TM-TM complexes and additional TM-soluble assemblies. Most predictions exhibited plausible membrane geometry: among 125,657 evaluable complexes with TM proteins possessing more than two TM helices, 95.4% had joint hydrophobic thicknesses within the expected range of 15-35 angstroms. Nevertheless, some complexes displayed substantial inconsistencies in membrane positioning or protomer orientation despite favourable prediction confidence. Interactions between TM proteins, particularly those containing only one or two TM helices, were frequently dominated by extramembrane contacts. Among TM-soluble complexes, 77.4% showed no penetration of the soluble partner into the hydrophobic membrane core. We further investigated whether AlphaFold3 (AF3) could overcome limitations of AlphaFold-Multimer. Although AF3 recovered an experimentally supported SERCA-phospholamban interaction, it failed to reproduce the cholesterol-mediated norepinephrine transporter homodimer. A benchmark of 173 cholesterol-containing membrane protein targets further revealed limited accuracy in predicting cholesterol-binding poses, particularly for structures released after the AF3 training cutoff. Together, these findings demonstrate that membrane-specific geometric constraints provide a valuable complement to conventional confidence metrics and highlight the persistent challenges of predicting assemblies involving lipids.

## Introduction

Protein–protein interactions (PPIs) are fundamental to nearly all cellular processes, including signal transduction, transport, metabolism, and the regulation of protein activity. Proteins frequently function as components of multimeric assemblies, in which interactions between individual subunits determine their localization, stability, and biological activity. Consequently, identifying interacting proteins and characterizing the molecular organization of their complexes are essential for understanding cellular function and disease mechanisms (Greenblatt et al. 2024). Considerable efforts have been devoted to mapping protein interaction networks using high-throughput experimental approaches, including yeast two-hybrid assays and affinity purification coupled with mass spectrometry. Large-scale studies, such as the Human Reference Interactome (HuRI) and BioPlex, have identified tens of thousands of protein interactions (Luck et al. 2020; Huttlin et al. 2021). These experimental datasets, together with interactions curated from the scientific literature and computationally inferred associations, are integrated into public resources such as IntAct, BioGRID, and STRING (Del Toro et al. 2022; Oughtred et al. 2021; Szklarczyk et al. 2025).

Although these resources provide extensive information on protein associations, they generally lack structural details and may include indirect interactions. Three-dimensional structures are therefore essential for identifying interaction interfaces, understanding molecular mechanisms, and interpreting disease-associated mutations. Experimentally determined structures of protein assemblies are available through the Protein Data Bank (PDB), primarily obtained using X-ray crystallography, cryo-electron microscopy, and nuclear magnetic resonance spectroscopy (wwPDB consortium 2019). However, despite substantial progress in experimental structural biology, the number of characterized protein complexes remains limited relative to the large number of known and predicted protein interactions. This limitation is particularly pronounced for transmembrane (TM) proteins, whose experimental characterization is complicated by their hydrophobic surfaces, conformational flexibility, and dependence on the surrounding lipid environment (Hegedűs et al. 2022; Lin et al. 2026; Q. Zhang and Cherezov 2019).

Computational approaches have therefore become increasingly important for determining the structures of protein complexes. Earlier methods primarily relied on structural templates or molecular docking, using geometrical complementarity, physicochemical interactions, and experimentally derived restraints to predict the relative orientations of interacting proteins. Widely used examples include HADDOCK and ClusPro (Dominguez et al. 2003; Kozakov et al. 2017). More recently, advances in deep learning have substantially transformed protein structure prediction. AlphaFold2 provided highly accurate predictions of individual protein structures, while RoseTTAFold and AlphaFold-Multimer extended these capabilities to protein assemblies (Jumper et al. 2021; Evans et al. 2021; Baek et al. 2021). Early studies demonstrated the feasibility of AlphaFold2-based protein complex prediction and extended its application to large-scale modeling of the human protein interactome (Bryant et al. 2022; Burke et al. 2023). AlphaFold3 further expanded structural prediction to complexes containing proteins, nucleic acids, small molecules, and other molecular components (Abramson et al. 2024). These advances have enabled large-scale exploration of the structural interactome. More recently, ESMFold2 has demonstrated accurate protein-complex prediction using protein language model representations, without requiring multiple sequence alignments (Candido et al. 2026). In parallel, the AlphaFold Protein Structure Database (AFDB) was extended to include approximately 31 million candidate homo- and heterodimeric complexes predicted using AlphaFold-Multimer, of which 1.81 million satisfied stringent interface-confidence criteria (Han et al. 2026). Among these predicted assemblies, transmembrane (TM) protein complexes represent a particularly interesting class, as their structures and interactions are constrained by the physicochemical properties of the lipid bilayer.

Although AlphaFold-based approaches have demonstrated considerable accuracy in predicting TM protein structures (Hegedűs et al. 2022), standard protein-complex predictions do not explicitly account for the surrounding lipid bilayer. This limitation may affect the predicted orientation and association of membrane proteins, particularly when interactions involve extramembrane domains or membrane lipids. However, the lipid bilayer also imposes physical constraints that can be exploited to assess prediction quality. Proteins embedded within the same membrane must adopt compatible membrane positions, orientations, and hydrophobic thicknesses, while soluble interaction partners should generally remain outside the hydrophobic bilayer core. These constraints provide an additional level of structural assessment complementary to conventional prediction-confidence metrics. In this study, we systematically evaluated the membrane compatibility of high-confidence AlphaFold-Multimer complex predictions available through AFDB, including TM–TM and TM–soluble complexes (Supplementary Figure S1). These geometric assessments revealed generally favorable membrane compatibility, while identifying a subset of complexes with substantial inconsistencies. We further investigated whether AlphaFold3 could resolve some of these inconsistencies and evaluated its ability to reproduce experimentally observed cholesterol-mediated protein interactions.

## Results

### Interface composition of transmembrane protein complexes

We first examined the location of predicted protein–protein interfaces relative to the membrane in high-confidence AFDB complexes containing at least one transmembrane helix in each protomer. The dataset comprised 6,027 heterodimeric and 183,442 homodimeric complexes, totaling 189,469 TM–TM complexes. TM helices were assigned using TMbed (Bernhofer and Rost 2022), and inter-protein contacts were classified according to whether they involved TM or soluble regions. In most complexes, contacts involving soluble regions predominated, whereas interfaces dominated by TM–TM contacts represented a smaller fraction of the dataset (Figure 1A). This distribution is expected because TM helices typically contain only approximately 20–25 residues and therefore provide relatively limited interaction surfaces, whereas many membrane proteins contain substantially larger soluble domains capable of forming extensive interfaces.

**Figure 1.**
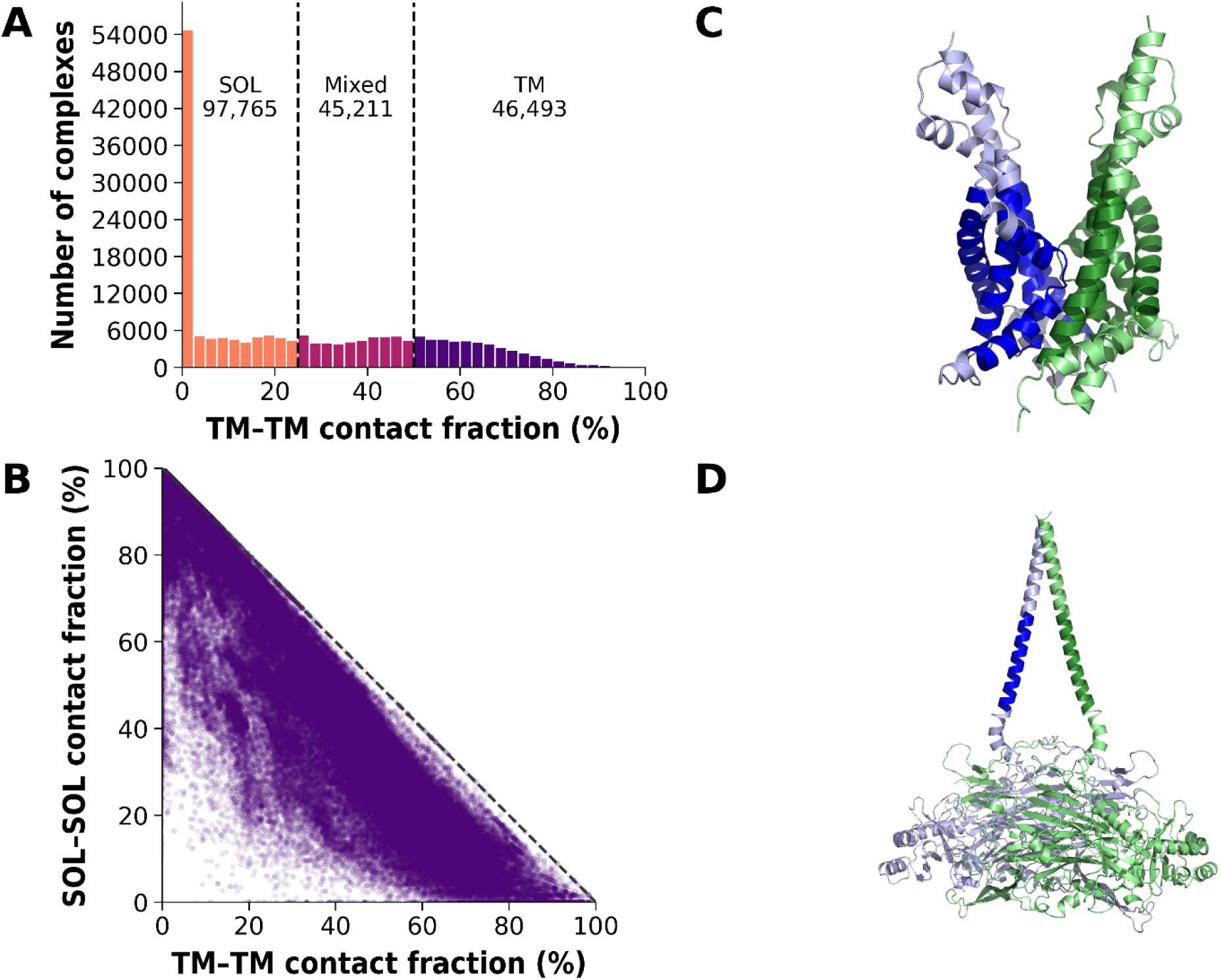
Interface types of transmembrane protein dimers. **(A)** Distribution of TM–TM contact fractions across the dataset (N = 189,469), with SOL-, Mixed-, and TM-interface classes separated by the 25% and 50% thresholds (dashed vertical lines). **(B)** Relationship between the SOL–SOL contact fraction and the TM–TM contact fraction for individual complexes. Each point represents one complex. The dashed diagonal line indicates the theoretical boundary at which TM–TM fraction + SOL–SOL fraction = 100%; points below this line therefore correspond to complexes with a non-zero fraction of mixed TM–SOL contacts. **(C)** A representative example of a TM-interface complex (AF-0000000070197049), in which the interaction is dominated by contacts between transmembrane helices. **(D)** A representative example of a SOL-interface complex (AF-0000000082824603), in which the interaction is dominated by contacts between soluble regions. TM regions are indicated by darker colors.

To examine the relationship between membrane-embedded and soluble interactions more directly, the SOL–SOL contact fraction was plotted against the TM–TM contact fraction for each complex (Figure 1B). This representation reveals the expected trade-off between these two interface components while also allowing complexes containing substantial mixed TM–SOL contacts to be distinguished. Representative examples with high TM–TM contact fractions and with predominantly SOL–SOL contacts are shown in Figure 1C and D, respectively.

### Membrane geometry and compatibility of highTM–highTM complexes

To assess membrane geometry, we focused on complexes in which both protomers contained at least three TM helices (highTM– highTM). Three or more TM helices provide sufficient spatial information to estimate membrane orientation and boundaries independently for each protomer. Membrane geometry could be evaluated for 125,657 complexes, including 122,553 homodimers and 3,104 heterodimers, with at least three structurally usable TM helices per protomer.

We first assessed membrane compatibility by calculating the joint hydrophobic thickness of each highTM–highTM complex from the pooled intracellular and extracellular TM-terminal coordinates of both protomers. This complex-level measurement reflects their collective membrane-spanning geometry and is sensitive to differences in the orientations of their TM regions. The resulting distribution was concentrated within the expected hydrophobic-thickness range of 15–35 Å, with 95.4% of the 125,657 complexes falling within this interval and a median thickness of 24.0 Å (Figure 2A). Nevertheless, a reduced joint thickness may reflect either atypical individual TM geometry or disagreement between protomer orientations. The homodimeric structure of the E3 ubiquitin-protein ligase RNFT1 is an outlier example with extreme geometric deviation and is presented in Figure 2B. Although several membrane-topology predictors broadly agree on the locations of the transmembrane regions, these assignments remain somewhat uncertain and only loosely define the expected bilayer boundaries. More importantly, the two protomers adopt markedly different orientations, such that the membrane planes inferred from the individual protomers are approximately perpendicular rather than parallel. This example therefore suggests a potential AlphaFold failure despite otherwise favorable confidence scores (Supplementary Figure S2). In contrast, other highly suspicious complexes could be explained more directly by incorrect transmembrane-topology predictions or annotations (Supplementary Figure S3).

**Figure 2.**
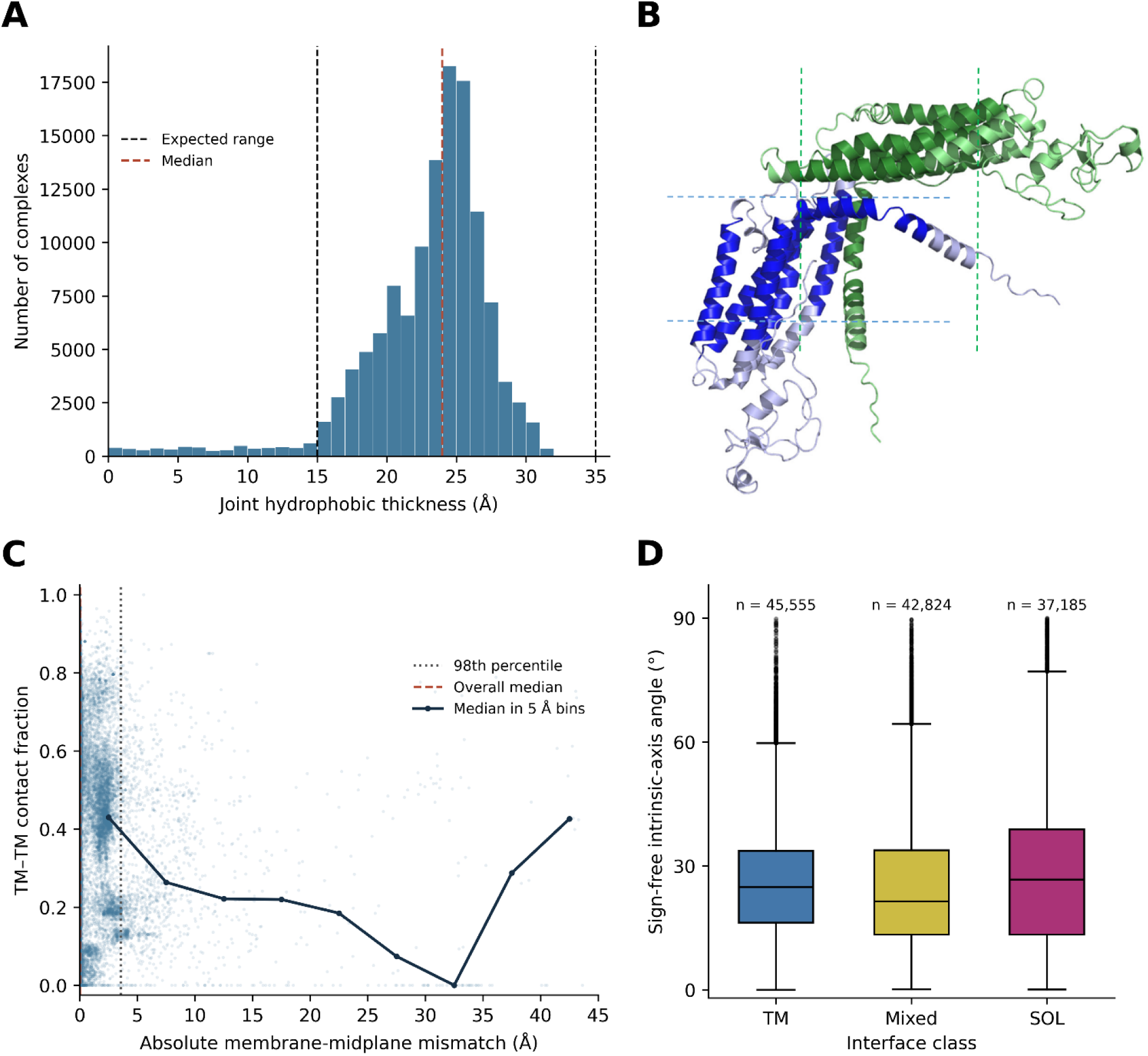
Bilayer geometry and compatibility of highTM–highTM protein complexes. **(A)** Distribution of hydrophobic bilayer thickness derived from the transmembrane geometry of highTM protein dimers. Dashed vertical lines indicate the expected hydrophobic-thickness range based on experimental membrane-protein structures, and the solid line indicates the median. **(B)** The homodimeric structure of the E3 ubiquitin-protein ligase RNFT1, exhibits valid bilayer thickness, but protomer orientation is likely non-physiological. Protomers are colored blue and green. TM regions are indicated by darker colors. Lipid bilayers are indicated by dashed lines to illustrate that the two protomers are oriented approximately perpendicular to each other, enclosing an angle of ∼90°, rather than adopting the expected near-parallel orientation. **(C)** TM–TM contact fraction as a function of the absolute membrane-midplane mismatch between interacting protomers. The vertical dashed line indicates the 98th percentile and the dash-dotted curve the median calculated in 5 Å mismatch bins. For visual clarity, 25 complexes with membrane-midplane mismatches greater than 45 Å are not displayed. **(D)** Distribution of the sign-free intrinsic-axis angle between protomers for TM-dominated, Mixed, and SOL-dominated interfaces. Smaller angles indicate more closely aligned membrane orientations.

Conversely, an apparently plausible joint thickness does not necessarily indicate that the two protomers occupy compatible membrane positions. We therefore examined the relative membrane placement and orientation of the protomers separately. This analysis included 80,089 complexes with relative TM-axis angles ≤30° and available inter-protein contact fractions, thereby restricting the comparison to protomers with sufficiently similar membrane orientations. The membrane-midplane mismatch describes the absolute displacement between the membrane midplanes defined by the two protomers along their common membrane-normal direction (Supplementary Figure S1). Thus, values close to zero indicate that both proteins independently place the membrane at approximately the same position, whereas large values indicate incompatible membrane placement. In the combined homo- and heterodimer dataset, the mismatch distribution was strongly concentrated at small values, with the 98^th^ percentile at 3.6 Å (Figure 2C). TM–TM contact fractions tended to be higher at small membrane-midplane mismatches, whereas the limited number of complexes with large mismatches prevented a reliable assessment of the relationship in this range.

Analysis of homo- and heterodimers separately revealed a substantially broader distribution among predicted heterodimers (Supplementary Figure S4). Whereas homodimers showed close agreement between the membrane positions derived from their two protomers, the 95th percentile of the heterodimer distribution reached approximately 10.5 Å. This difference is expected in part because identical protomers inherently have similar membrane geometry, whereas different proteins may prefer different hydrophobic thicknesses and membrane positions.

Finally, we examined the relative orientation of the protomers using the sign-free intrinsic-axis angle (Figure 2D). Of the 125,657 complexes with evaluable membrane geometry, 125,564 had assigned interface classes and were included in the comparison of relative TM-axis angles among interface types. This quantity describes the angle between the membrane axes independently derived from the two proteins, irrespective of axis direction (values close to 0° correspond to nearly parallel membrane orientations). The distributions slightly differed according to interface type. SOL-dominated complexes showed the highest median angle and the broadest distribution, indicating substantially greater freedom in the relative orientation of their transmembrane regions. TM-dominated complexes exhibited a lower median angle and a narrower distribution, while mixed interfaces showed the lowest median angle. These results indicate that direct TM–TM association imposes stronger geometrical constraints on the relative orientation of the interacting membrane proteins than interactions mediated predominantly by their soluble regions.

### Extending Membrane-Geometry Assessment to Single- and Double-Pass TM Protein Complexes

We next analyzed complexes between lowTM proteins containing one or two transmembrane helices and highTM proteins containing at least three usable TM helices. Because the highTM partner provides a more robust definition of membrane orientation and hydrophobic thickness, it was used as the reference for assessing the geometry of the lowTM partner. After restricting the analysis to complexes with an evaluable highTM reference thickness of 15–35 Å, 1,356 complexes remained, including 710 1TM– highTM and 646 2TM–highTM complexes. Comparison of the projected hydrophobic thickness of the lowTM partner with the membrane thickness derived from the highTM partner showed that most complexes were compatible with placement in a common membrane (Figure 3A, Supplementary Figure S5). Overall, 905/1,356 (66.7%) complexes were classified as geometrically compatible, 242 (17.8%) as borderline, and 209 (15.4%) as suspicious. Compatibility was substantially higher among 2TM–highTM complexes (505/646; 78.2%) than among 1TM–highTM complexes (400/710; 56.3%).

**Figure 3.**
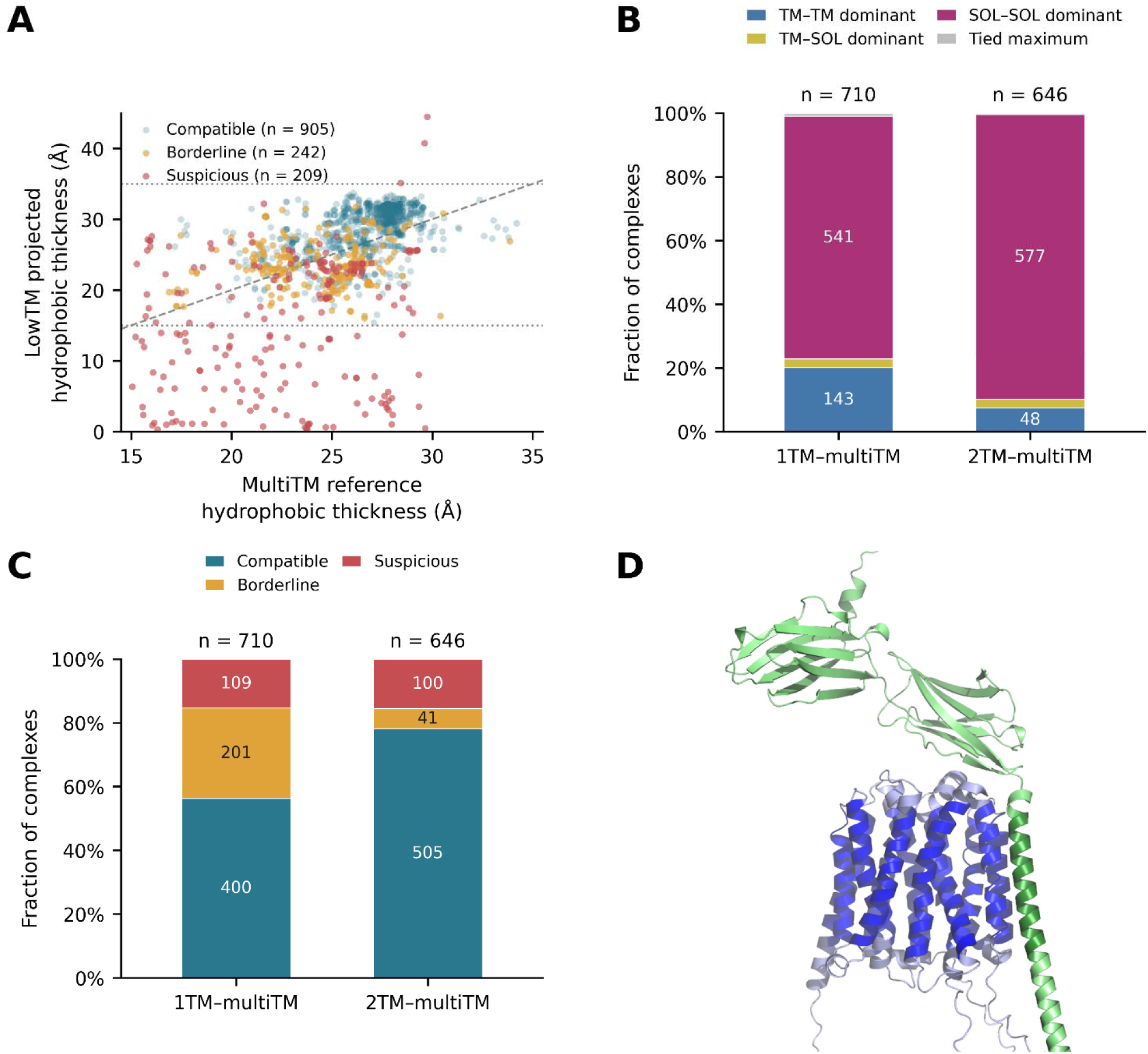
Interface composition and membrane-geometry compatibility of lowTM–highTM complexes. **(A)** Comparison of the hydrophobic thickness of the highTM reference partner with the projected hydrophobic thickness of the lowTM partner for the QC-eligible set of 1,356 complexes. Points are colored according to the overall geometry classification as Compatible, Borderline, or Suspicious. The dashed diagonal indicates equal thickness, whereas horizontal guidelines at 15 and 35 Å delimit the accepted projected hydrophobic-thickness range for the lowTM partner. **(B)** Interface composition of lowTM–highTM complexes. Bars show the fractions of complexes in which TM–TM, TM–SOL, or SOL–SOL contacts constitute the largest fraction of inter-protein contacts; exact ties are shown separately. **(C)** Overall geometry classification for lowTM–highTM complexes and for the combined cohort. **(D)** Representative lowTM–highTM complex illustrating a geometrically compatible arrangement of the lowTM partner (embigin) within the membrane defined by the highTM reference (Monocarboxylate transporter 3; AF_0000000210521500).

Despite containing membrane-spanning segments, most lowTM–highTM interfaces were dominated by contacts outside the TM regions (Figure 3B). SOL–SOL contacts were dominant in 541/710 (76.2%) 1TM–highTM complexes and 577/646 (89.3%) 2TM–highTM complexes. TM–TM-dominant interfaces were considerably less frequent, accounting for 143/710 (20.1%) and 48/646 (7.4%), respectively, while TM–SOL-dominant interfaces represented only ∼3% of either group. Thus, even when both partners are integral membrane proteins, association is frequently driven primarily by their extramembrane regions rather than by direct TM-helix packing.

Notably, 2TM proteins showed a higher fraction of geometrically compatible complex arrangements than 1TM proteins (Figure 3C). This likely reflects the greater orientational and positional constraints imposed by two membrane-spanning helices, which more strongly restrict the placement of the lowTM partner than a single TM helix. A representative geometrically compatible lowTM–highTM complex illustrates how the lowTM partner can be positioned consistently within the membrane defined by the highTM protein (Figure 3D).

Extending the analysis to lowTM–lowTM complexes required an empirical reference framework, since neither protomer contains enough TM helices to independently define membrane boundary planes. We therefore constructed topology-specific empirical references from 1,587 geometrically compatible (TM-axis angle ≤40° and minimum individual hydrophobic thickness ≥15 Å) highTM–highTM complexes. One or two TM helices were sampled from each protomer to generate pseudo-1TM–1TM, pseudo-1TM–2TM, and pseudo-2TM–2TM representations (Supplementary Figure S6A). Repeated sampling was used to characterize the expected distributions of midplane mismatch, relative TM-axis angle, and projected TM-span difference (Supplementary Figure S6B–C). An inferred membrane thickness, defined as the smaller projected TM span of the two partners, was additionally evaluated against the 15–35 Å membrane-spanning interval (Supplementary Figure S6E).

The lowTM–lowTM dataset contained 610 complexes, comprising 459 1TM–1TM, 62 1TM– 2TM, and 89 2TM–2TM complexes (Figure 4A). Geometry could be evaluated for 540 complexes. 447 were classified as geometrically consistent, 41 as suspicious, and 52 as highly suspicious, while 70 remained unevaluable. The largest fraction of highly suspicious arrangements occurred among 2TM– 2TM complexes, with 20 of 82 evaluable cases (24.4%), compared with 27 of 404 1TM–1TM and 5 of 54 1TM–2TM complexes. Despite both partners containing TM helices, their predicted interfaces were overwhelmingly dominated by contacts outside the annotated TM regions. SOL–SOL contacts were dominant in 593 of 610 complexes (97.2%), whereas only 15 complexes were TM–TM dominant and two were TM–SOL dominant (Figure 4B). All TM–TM-dominant cases occurred in the 2TM–2TM group. Moreover, among geometrically consistent 2TM–2TM complexes only 1 of 59 was TM–TM dominant, whereas 14 of 20 highly suspicious complexes showed TM–TM-dominated interfaces (Figure 4C). Thus, TM–TM-dominated interactions are rare among predicted lowTM–lowTM complexes that display membrane-compatible geometry, while most predicted interactions are instead mediated predominantly by regions outside the annotated TM segments.

**Figure 4.**
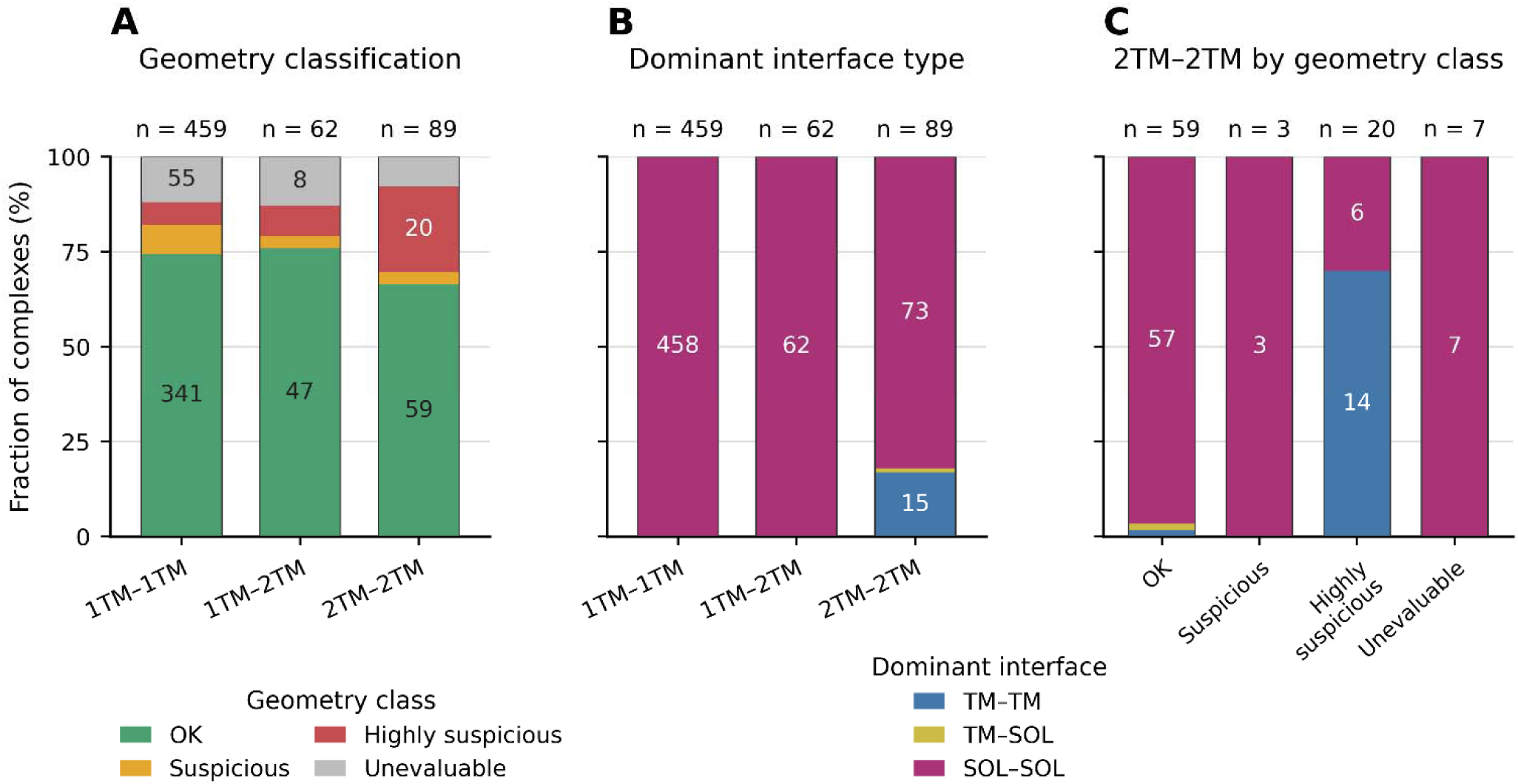
Geometry and interface composition of lowTM–lowTM complexes. **(A)** Geometry classification of 1TM–1TM, 1TM–2TM, and 2TM–2TM complexes using the topology-specific empirical reference derived from compatible highTM–highTM complexes. Complexes are classified as geometrically consistent, suspicious, highly suspicious, or unevaluable. The total number of complexes in each topology is indicated above the bars. **(B)** Dominant interface type for the same lowTM–lowTM complexes. Interface dominance was defined as the largest of the TM–TM, TM–SOL, and SOL–SOL contact fractions; no >50% threshold was required. SOL–SOL contacts dominate almost all 1TM–1TM and 1TM–2TM complexes and most 2TM–2TM complexes. **(C)** Dominant interface type of 2TM–2TM complexes stratified by geometry class. TM–TM-dominated interfaces are strongly concentrated among highly suspicious complexes, whereas geometrically consistent complexes are predominantly SOL–SOL dominated. Geometry-unevaluable complexes were retained in the interface analysis.

### Soluble partners generally remain outside the membrane core in highTM–soluble heterodimers

To assess the membrane compatibility of complexes between transmembrane and soluble proteins, we identified 4,643 heterodimers containing one highTM partner with at least three annotated TM helices and one partner without annotated TM segments. Sequence-based chain assignment was obtained for 4,642 complexes, and membrane geometry could be evaluated for all of these. After restricting the analysis to complexes with the TM-defined membrane thickness of 15–35 Å, 4,618 complexes were retained. In most complexes, the soluble partner remained outside the membrane-associated regions (Figure 5A). No soluble-partner residue entered the TM-defined hydrophobic core in 3,575 complexes (77.4%), while 2,598 complexes (56.3%) contained no residues in the 8 Å membrane-interface region. Accordingly, the median hydrophobic-core occupancy was zero, and even the 95th percentile corresponded to only 7.49% of the soluble-partner residues. Thus, extensive core occupation was restricted to a small subset of complexes.

**Figure 5.**
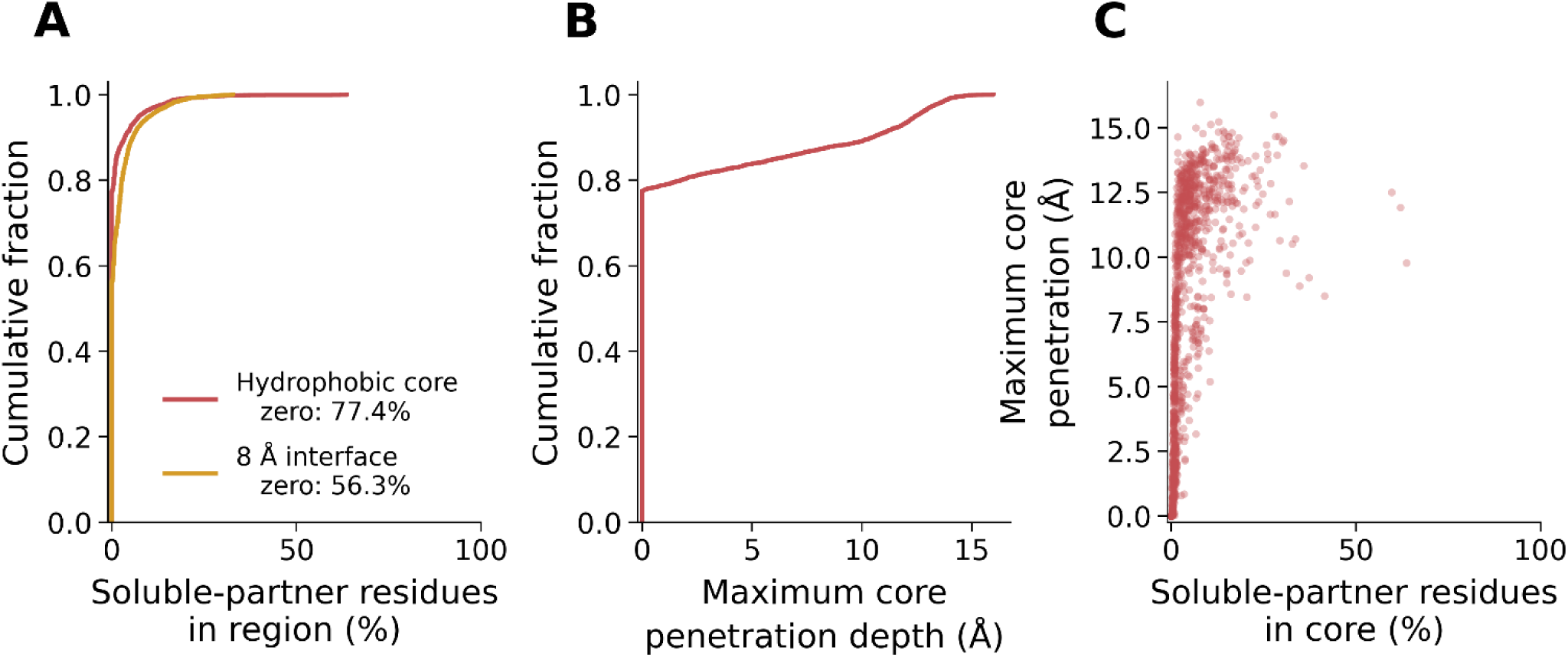
Membrane positioning of soluble partners in highTM–soluble heterodimers. Membrane-boundary planes were determined from the highTM partner, and complexes with a TM-defined membrane thickness of 15–35 Å were retained (*N* = 4,618). **(A)** Empirical cumulative distributions of the percentage of soluble-partner residues located in the hydrophobic core and interface. No core residues were observed in 77.4% of complexes, while 56.3% contained no residues in the interface. **(B)** Empirical cumulative distribution of the maximum penetration depth of soluble-partner residues into the hydrophobic core. Complexes without core penetration were assigned a depth of 0 Å. **(C)** Relationship between the percentage of soluble-partner residues located in the hydrophobic core and the maximum penetration depth. Each point represents one complex.

Maximum penetration into the TM-defined core showed a similarly strongly skewed distribution (Figure 5B). Because non-penetrating complexes were assigned a depth of zero, the median penetration depth was 0 Å, whereas the 95th percentile was 12.44 Å. Among the 1,043 complexes with at least one core residue, the median maximum penetration was 9.61 Å and the maximum was 15.97 Å.

Comparison of core occupancy with penetration depth revealed substantial variation among the core-positive complexes (Figure 5C). Some soluble partners penetrated relatively deeply while only a small fraction of their residues entered the membrane core, whereas extensive core occupancy was observed only in a limited number of complexes. Together, these results indicate that most predicted highTM–soluble heterodimers are geometrically compatible with placement of the soluble partner outside the hydrophobic membrane region, although a minority show substantial membrane-core overlap. Importantly, in some cases, disordered and highly dynamic regions of the soluble partner may extend across the inferred bilayer region and thereby contribute to geometrically less compatible predictions. This is not unexpected, because AlphaFold does not explicitly model the membrane bilayer, and such cases can be readily identified by geometric analysis (Hegedűs et al. 2022; Dobson et al. 20 3).

### AF3 improves the prediction of transmembrane protein interactions but remains limited in modeling lipid-mediated complexes

Complexes of SERCA, which are studied in our wet lab (Berta et al. 2024, 2026), with its known regulatory proteins were absent from the analyzed AlphaFold database of predicted protein complexes. To examine whether these interactions could be recovered by structure prediction, we modeled human SERCA2b in complex with phospholamban (PLN), a single-pass transmembrane protein that regulates SERCA activity. Initial predictions performed with AlphaFold-Multimer (AF2-Multimer) using default parameters failed to reproduce the expected interaction. None of the generated models exhibited direct contacts between SERCA and PLN. Moreover, PLN was erroneously positioned with its cytosolic N-terminal region facing the luminal side of SERCA, resulting in an incorrect relative membrane orientation (Figure 6A). In contrast, AF3 predicted a compact SERCA–PLN complex in which the transmembrane helix of PLN formed close contacts with the transmembrane region of SERCA (Figure 6B). This arrangement was consistent with the experimentally determined structure of rabbit SERCA1a in complex with PLN (PDB: 4y3u; Figure 6C). These results illustrate the improved ability of AF3 to recover a biologically relevant transmembrane protein interaction that was not reproduced by AF2-Multimer under the tested conditions.

**Figure 6.**
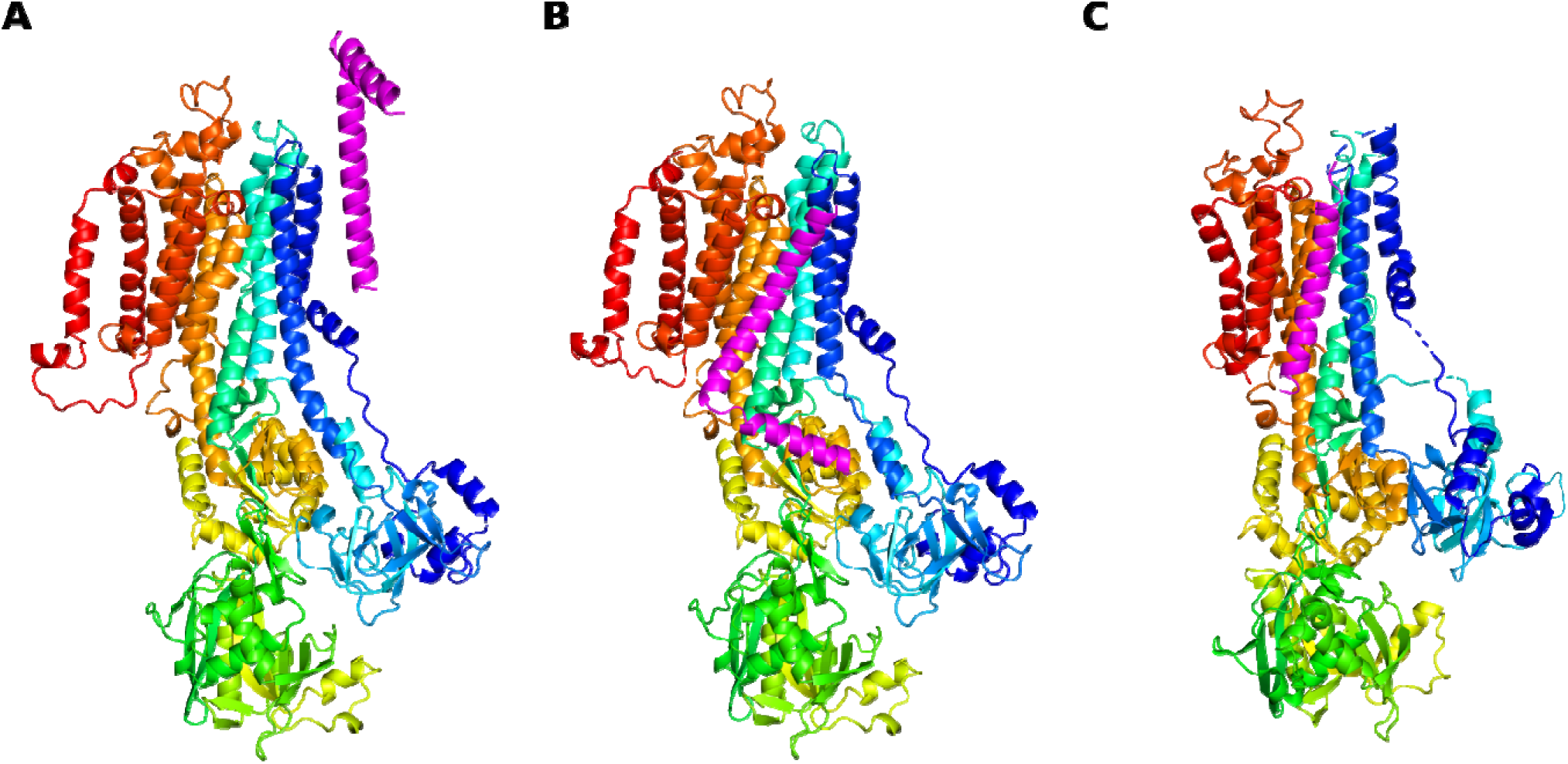
Comparison of AlphaFold-Multimer and AlphaFold 3 predictions of the SERCA– phospholamban complex. **(A)** AlphaFold-Multimer prediction showing an incorrect relative orientation of SERCA2b and phospholamban (PLN), with no direct contacts between the proteins. **(B)** AlphaFold3 prediction showing PLN interacting with the transmembrane region of SERCA2b in an arrangement consistent with the experimentally determined complex. **(C)** Experimental structure of rabbit SERCA1a in complex with PLN (PDB: 4y3u). In all panels, SERCA is colored from blue (N-terminus) to red (C-terminus), and PLN is shown in magenta.

Unlike AF2-Multimer, AF3 can explicitly incorporate non-protein components, including lipids and other small molecules, into complex structure predictions. This extends its potential applicability to membrane protein assemblies in which lipids contribute directly to protein–protein interactions. Examples of such assemblies include the urea transporter DUR3 (PDB: 9j7d), the mechanosensitive ion channel OSCA1.2 (PDB: 8t56), and the norepinephrine transporter NET (PDB: 9ke3) (An et al. 2025; Jojoa-Cruz et al. 2024; H. Zhang et al. 2025). We investigated NET in greater detail because its lipid-mediated homodimer is functionally relevant and, unlike OSCA1.2, its protomers are separated by lipids without direct protein–protein contacts. Moreover, cholesterol molecules occupy the interface between the NET protomers (Figure 7A). Cholesterol represents a comparatively favorable test case for AF3 because it is smaller and conformationally less flexible than phospholipids. AF3 predictions were performed locally using standard parameters with 14 cholesterol and two POPG molecules included. To increase sampling, 100 NET dimer structures were predicted, but none reproduce successfully the interface observed in the cryo-EM structure (Figure 7B,C).

**Figure 7.**
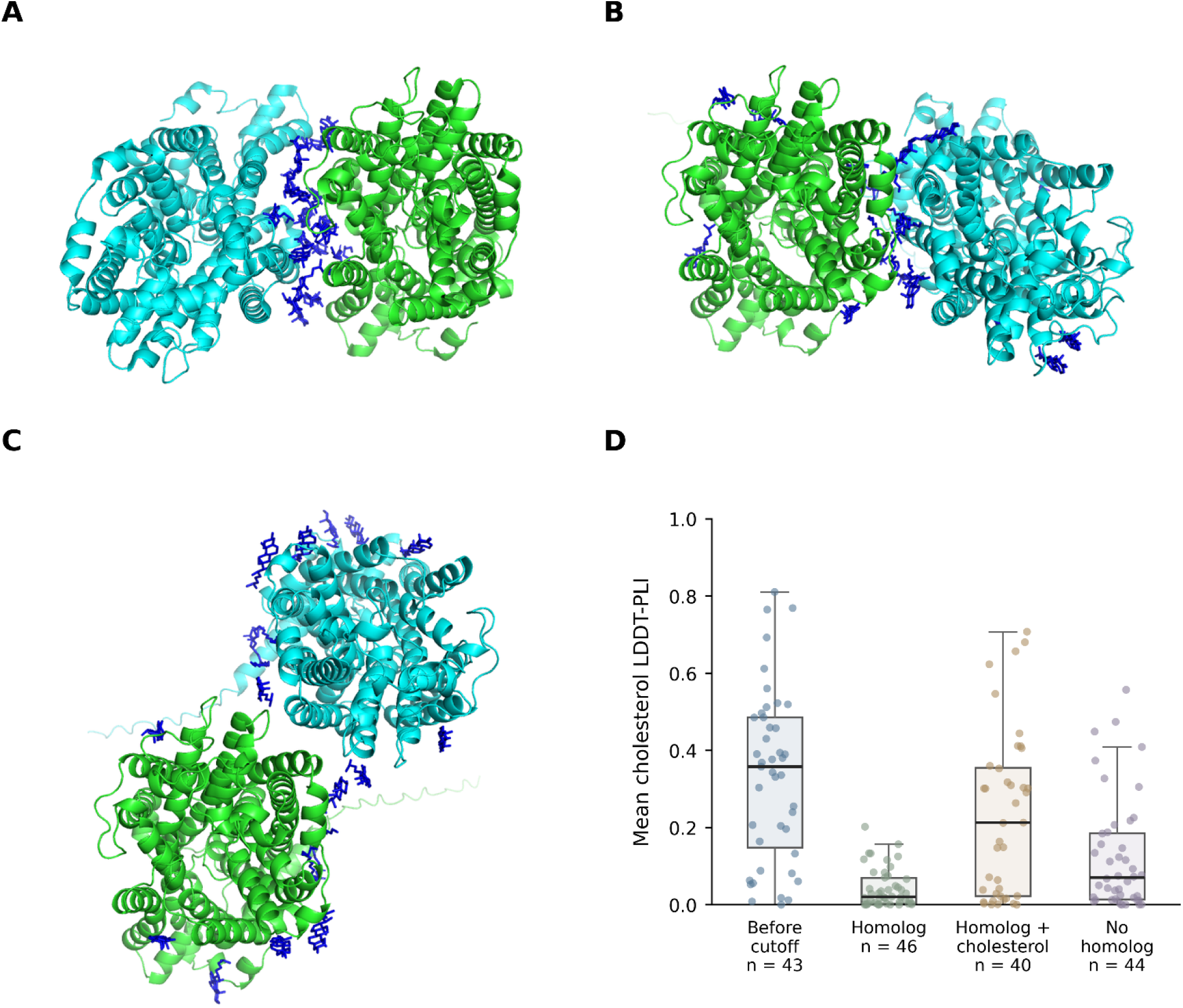
AF3 poorly reproduces the cholesterol-mediated NET dimer interface and cholesterol-binding sites. **(A)** Experimental cryo-EM structure of the NET dimer (PDB ID: 9ke3). The two protomers are shown in green and turquoise, and cholesterol molecules in blue. **(B, C)** Representative AF3 predictions illustrating the two distinct dimer interfaces observed among the 100 generated models. For comparison with the experimental structure, one predicted protomer (green) was aligned to the corresponding protomer of the cryo-EM structure. Neither predicted interface reproduces the experimentally observed cholesterol-mediated dimer arrangement. **(D)** Target-level distribution of mean cholesterol LDDT-PLI scores across the analyzed AF3 prediction groups, showing generally poor and inconsistent reproduction of experimentally observed cholesterol-binding poses.

The failure to reproduce the NET dimer could arise not only from inaccurate protein–protein assembly but also from limitations in predicting cholesterol-binding poses. To investigate the latter possibility systematically, we assembled a benchmark of 173 evaluable cholesterol-containing membrane protein targets. Targets were divided into four approximately equally sized groups according to the AF3 training cutoff and the availability of homologous structural information. Structures released before the 30 September 2021 cutoff formed the before-cutoff group, whereas post-cutoff targets were classified as homolog when a related structure without cholesterol was available, homolog+cholesterol when a cholesterol-containing homologous structure was available, or no homolog when no qualifying homologous structure was identified. The four groups contained 43, 46, 40, and 44 targets, respectively. AF3 showed limited accuracy in reproducing experimentally observed cholesterol-binding poses, with low mean target-level LDDT-PLI (Local Distance Difference Test for Protein–Ligand Interactions) scores across all three post-cutoff groups (0.043, 0.230, and 0.125 for homolog, homolog+cholesterol, and no-homolog targets, respectively; Figure 7D). The availability of a homologous structure without cholesterol was not associated with improved prediction accuracy, whereas cholesterol-containing homologs were associated with higher, although still relatively low scores.

## Discussion

The expansion of the AlphaFold Protein Structure Database (AFDB) to multimeric predictions offers substantial opportunities for studying protein interactions, but high interface confidence should not be equated with physiological relevance. The stringent selection of high-confidence complexes, including the use of ipSAE to reduce the influence of disordered or non-interacting regions, improves the interpretability of predicted interfaces without establishing that the interactions occur in cells (Han et al. 2026; Dunbrack 2025). The proportion of candidate models passing these filters is not itself a measure of predictive accuracy: some candidate pairs may not interact physiologically, and exclusion also reflects the chosen confidence thresholds. Studies of oligomeric-state prediction similarly emphasize the importance of structural context and the limitations of confidence-based inference, particularly for homodimers (Lin et al. 2026; Danielsson and Elofsson 2025). This distinction is especially important in the AFDB extension, which is dominated by homodimer predictions. Independent criteria are therefore needed to assess the biological plausibility of these assemblies.

For transmembrane (TM) complexes, the lipid bilayer provides particularly informative physical constraints. Most high-confidence AFDB predictions examined here showed plausible hydrophobic thicknesses and relative membrane arrangements, despite the absence of an explicit membrane during prediction. However, a subset exhibited substantial midplane displacement, incompatible orientations, or atypical thicknesses. The RNFT1 homodimer illustrates how a confident interface can coexist with questionable membrane geometry, although uncertainty in its TM annotation complicates interpretation (Supplementary Figure S2). Importantly, this is a complementary rather than redundant test. Confidence concerns relative structural uncertainty, whereas the geometry check evaluates whether two protomers can be placed coherently in a shared bilayer. Membrane geometry can thus flag predictions for further assessment, but geometric compatibility alone cannot validate an interaction or establish its physiological oligomeric state. Moreover, the overall distributions are dominated by homodimers, and the broader geometry observed for heterodimers cautions against extrapolating pooled results to all interaction types.

A related finding was the predominance of extramembrane contacts, especially in complexes containing one or two TM helices. Such interfaces can be physiologically reasonable because TM segments may act mainly as membrane anchors. Nevertheless, confident packing between soluble domains does not guarantee compatible positioning of the membrane-spanning regions. The enrichment of TM–TM-dominated interfaces among highly suspicious 2TM–2TM predictions is particularly informative, as seemingly extensive TM packing can arise from an assembly that violates membrane constraints. This reinforces the need to consider interface composition and membrane geometry together, rather than treating either as sufficient evidence of a correct assembly.

Most predicted TM–soluble heterodimers also positioned the soluble partner outside the hydrophobic membrane core. Yet apparent penetration should be interpreted cautiously: amphipathic helices, lipid-modified regions, partially inserted elements, and intrinsically disordered segments can overlap the geometrically inferred membrane region without demonstrating an incorrect protein–protein interaction (Hegedűs et al. 2022; Dobson et al. 2023). Such cases require inspection of the implicated residues and their structural confidence rather than automatic rejection. In particular, penetration caused by an uncertain, flexible terminal segment should not be weighted like penetration by a compact, well-resolved soluble domain.

Interactions with small, single-pass regulatory proteins present a further challenge because their interfaces may be limited or conformationally dependent. Although some accessory membrane-protein interactions are represented in the predicted dataset, established SERCA regulatory complexes were absent. In our test case, AlphaFold-Multimer did not recover the SERCA–phospholamban association, whereas AF3 produced a membrane-compatible arrangement resembling the experimental complex (Akin et al. 2013). This demonstrates that an interaction absent from the filtered AFDB, or missed in an initial AlphaFold-Multimer run, may still be recoverable with another prediction strategy. A single example, however, cannot establish a general improvement in AF3 accuracy across regulatory TM complexes, particularly when the possible roles of templates and conformational states have not been systematically compared.

AF3 also extends complex prediction to non-protein components, including lipids that may stabilize membrane proteins or mediate associations without direct protein–protein contacts (Abramson et al. 2024). The cholesterol-mediated norepinephrine transporter (NET) homodimer provided a stringent test of this capability, since its protomers are separated by lipids rather than a direct protein interface, and cholesterol is less conformationally flexible than typical phospholipids. Even with cholesterol and phospholipid molecules included, the additional AF3 sampling did not reproduce the experimentally observed dimer interface (Figure 7). This failure is notable because a correct lipid-mediated assembly requires both plausible lipid placement and the appropriate relative positioning of protein subunits. However, the present predictions cannot determine which source of error is primary.

Our cholesterol benchmark points to a related limitation and shows that AF3 generally reproduced post-training-cutoff cholesterol poses poorly. Cholesterol-containing homologous structures were associated with somewhat better predictions than homologs lacking cholesterol, but the comparison is observational and the accuracy remained limited. The independent LiPP benchmark likewise reported reduced success for previously unseen nonannular lipid poses and especially poor performance for annular lipids (Yang et al. 2026). Although its targets and evaluation criteria differ from ours, these findings support the conclusion that explicit inclusion of a lipid does not ensure an accurate protein–lipid interaction. Improved lipid placement may therefore be important for modeling lipid-mediated oligomers, but whether it would rescue the NET interface remains unresolved.

Several limitations qualify these conclusions. Geometry-based assessments depend on TM boundaries and on approximating the membrane as a common, locally planar bilayer; annotation errors, strong helix tilt, and unusual membrane architectures can create false warnings. The TAP1-containing models illustrate the distinction between geometrical classification and biological plausibility: assemblies without established physiological support must be distinguished from an authentic TAP1– TAP2 heterodimer whose apparent mismatch is sensitive to TM-region assignment (Supplementary Figure S3). Our analysis also tests plausibility rather than interaction validity, and the direct AlphaFold-Multimer–AF3 comparisons involve selected systems rather than a representative benchmark. Finally, experimentally observed cholesterol poses depend on protein conformation, ligand occupancy, and the local membrane environment, which were not exhaustively sampled here.

Together, these results support a layered evaluation of predicted membrane complexes in which confidence scores are complemented by TM topology, membrane geometry, interface composition, and biological evidence. Suspicious cases can be prioritized for alternative predictions, targeted sampling, membrane-aware refinement, or experimental testing. Importantly, the observed limitations do not diminish the substantial value of the expanded AFDB. Rather, they identify where additional information is needed to distinguish confident predictions from physiologically plausible molecular assemblies. Combining advances in multimeric structure prediction with membrane-specific physical constraints offers a promising route toward more reliable structural models of membrane protein interactions and, ultimately, a more accurate understanding of their mechanisms and functional consequences.

## Acknowledgments

We thank Daniel Nemeskeri for his assistance with the analysis of single-pass transmembrane proteins. We acknowledge the Digital Government Development and Project Management Ltd. for awarding us access to the Komondor HPC facility based in Hungary and the Scientific Computing Laboratory of the HUN-REN Wigner Research Centre for Physics for their support. This work was supported by the National Research, Development and Innovation Office of Hungary through grants TKP2021-EGA-24, NKFIH 152735, and HU-RIZONT-2024-00003.

## Data availability

Associated data are deposited at Zenodo (DOI:10.5281/zenodo.23940760) and GitHub (https://github.com/hegelab/AF-TM-Dimers-QC).

## Methods

### AlphaFold-Multimer complex datasets

Metadata for high-confidence homodimeric and heterodimeric AlphaFold-Multimer predictions were obtained from the European Bioinformatics Institute (https://ftp.ebi.ac.uk/pub/databases/alphafold/collaborations/nvda; (Han et al. 2026). Complexes containing at least one TM protein were identified using the TM annotation procedure described below, and their coordinates were retrieved using the AlphaFold Protein Structure Database (AFDB) API (Varadi et al. 2022). The numbers of complexes retained at successive stages of analysis, together with the corresponding eligibility criteria and figures, are summarized in Supplementary Table S1.

### TM protein identification and TM boundary detection

TM proteins and their membrane-spanning regions were identified using TMVisDB (Olenyi et al. 2025). Its TMbed-based annotations were used to assign residue-level TM regions (Bernhofer and Rost 2022). For complexes containing one identified TM protein and a partner without a TMVisDB annotation, TMbed was run locally to determine whether the unannotated partner contained TM segments. Of 3,244 analyzed partners, 137 were identified as TM proteins and 3,107 as non-TM, including proteins with signal-peptide-only predictions. TM annotations were mapped to the corresponding AFDB protomers using UniProt accessions.

### Inter-protein contact composition

For highTM–highTM dimers, the location of the protein–protein interface was quantified from interchain residue contacts. Two residues were considered to be in contact when at least one pair of non-hydrogen atoms, one from each residue, was within 5.0 Å. Each contacting residue pair was counted only once, irrespective of the number of atom-atom contacts between the residues. Complexes with fewer than 20 interchain contacting residue pairs were considered insufficiently supported for contact-composition classification.

Residues were assigned as transmembrane (TM) or non-TM according to the project TM annotations; TM residues corresponded to TM-helix (H/h) segments, whereas signal-peptide annotations were not considered TM segments. Interchain contacts were separated into TM–TM, TM–SOL, and SOL–SOL classes according to the annotation of the two contacting residues. The TM–TM contact fraction was calculated relative to all interchain contacting residue pairs. Interfaces were operationally classified as SOL-interface when fTM-TM<0.25, Mixed-interface when 0.25≤f_TM−TM_<0.50, and TM-interface when fTM−TM≥0.50. The 0.50 cutoff has a direct majority interpretation, identifying interfaces in which at least half of all inter-protein contacts are formed between TM residues. The 0.25 cutoff separates interfaces with only a minor TM contribution from those in which TM–TM contacts constitute a substantial component of the interface despite not being dominant. These thresholds were therefore used as operational boundaries for describing interface composition rather than as intrinsic biological discontinuities.

### Membrane-geometry assessment of highTM–highTM complexes

For complexes containing at least three usable TM helices per protomer, membrane geometry was determined independently for each protein. TM segments of at least 15 residues with available Cα coordinates at both termini were used to calculate membrane orientation. Each helix was represented by a normalized vector connecting its terminal Cα atoms, and the principal eigenvector of the sum of the outer products of these vectors defined the intrinsic membrane axis of each protomer. The relative orientation was quantified as the sign-independent angle between the two axes, ranging from 0° to 90°. To assess relative membrane positioning, a common reference axis was constructed by aligning the signs of the two protomer axes and normalizing their vector sum. The terminal coordinates of each TM helix were projected onto this axis, and the lower and upper membrane boundaries of each protomer were estimated as the medians of the corresponding projected helix endpoints. The membrane midpoint was defined as the average of these boundaries, and membrane-midplane mismatch was calculated as the absolute difference between the two protomer midpoints. Because this comparison becomes less reliable for strongly misaligned protomers, quantitative analysis of membrane-midplane mismatch was restricted to complexes with a relative axis angle ≤30°.

Hydrophobic thickness was calculated separately using the TM annotations to identify intracellular and extracellular helix termini. For each protomer, Cα coordinates from the first and last three residue positions of the annotated TM helices were pooled according to their membrane-side assignments. The Euclidean distance between the resulting intracellular and extracellular centroids defined the hydrophobic thickness. Values of 15–35 Å were considered consistent with the expected hydrophobic dimensions of biological membranes (Hegedűs et al., 2022).

### Assessment of membrane compatibility in lowTM–highTM complexes

Complexes containing one protein with one or two usable transmembrane helices (lowTM) and another with at least three usable TM helices (highTM) were analyzed using the highTM partner as the membrane reference. Side 1 and side 2 terminal centroids were calculated for each protomer by averaging pooled Cα coordinates from the first and last three residue positions of the annotated TM helices, with helix orientation used to assign the corresponding membrane sides. Helices lacking usable coordinates at either terminus were excluded.

The vector connecting the side 1 and side 2 centroids of the highTM partner defined the reference membrane normal, while its length defined the reference hydrophobic thickness. The geometric compatibility of the lowTM partner was evaluated using three measures: (i) projected hydrophobic thickness, calculated as the absolute projection of its side 1/side 2 centroid vector onto the reference membrane normal; (ii) the sign-independent angle between the side 1/side 2 vectors of the two partners; and (iii) the absolute displacement between the protomer midpoints projected onto the reference membrane normal.

Of 1,591 candidate complexes, 1,356 had a reference highTM thickness within the accepted 15– 35 Å interval and were retained for classification. Projected lowTM thicknesses within 15–35 Å were considered compatible, whereas values outside this range were suspicious. Relative angles were classified as compatible (≤40°), borderline (>40° to ≤50°), or suspicious (>50°). Similarly, midpoint displacements were classified as compatible (≤5 Å), borderline (>5 to ≤10 Å), or suspicious (>10 Å).

The overall classification was determined by the most severe individual criterion: complexes were classified as suspicious if any criterion was suspicious, borderline if at least one criterion was borderline and none was suspicious, and compatible only if all three criteria were compatible. The final dataset comprised 710 1TM–highTM and 646 2TM–highTM complexes.

### Empirical calibration of lowTM–lowTM membrane geometry

Because proteins containing only one or two TM helices cannot reliably define membrane boundaries independently, topology-specific reference distributions were constructed from 1,587 highTM–highTM complexes previously classified as geometrically compatible (relative TM-axis angle ≤40° and minimum individual hydrophobic thickness ≥15 Å). One or two TM helices of at least 15 residues were randomly selected from each protomer to generate pseudo-1TM–1TM, pseudo-1TM–2TM, and pseudo-2TM–2TM representations. For each source complex and topology, 100 random samples were generated, retaining the original structural coordinates.

The geometric descriptors included membrane-midplane mismatch, relative TM-axis angle, projected TM-span difference, and extracellular and intracellular boundary mismatches. To avoid pseudoreplication, topology-specific P95 and P99 reference thresholds were calculated from a separate fixed-seed selection containing one pseudo-complex per source complex and topology (N = 1,587 per topology). The repeated samples were used to characterize geometric variability, while only the independent fixed-seed selection was used to establish the classification thresholds.

### Analysis of real lowTM–lowTM complexes

Real lowTM–lowTM complexes were divided into three topology classes: 1TM–1TM, 1TM–2TM, and 2TM–2TM. TM helices were analyzed using the same boundary, midpoint, axis, orientation, and projection definitions as in the pseudo-lowTM calibration. For each complex, extracellular and intracellular boundary mismatches, membrane-midplane mismatch, relative TM-axis angle, projected TM spans, and projected span difference were calculated.

Geometric compatibility was evaluated using three descriptors: membrane-midplane mismatch, relative TM-axis angle, and projected TM-span difference. Each descriptor was compared with the P95 threshold of its corresponding topology-specific reference distribution. Complexes exceeding the P95 thresholds for at least two of the three descriptors were classified as suspicious. Complexes with fewer than two exceedances were considered geometrically consistent, unless they violated the additional hydrophobic-thickness criterion described below. For each complex, joint hydrophobic thickness was calculated by pooling the intracellular and extracellular TM-terminal Cα coordinates from both protomers, using the same three-residue terminal segments as for individual thickness calculations. The Euclidean distance between the resulting pooled intracellular and extracellular centroids defined the joint hydrophobic thickness. Values outside the inclusive 15–35 Å interval were classified as highly suspicious, irrespective of the percentile-based descriptors, overriding any other classification (Hegedűs et al. 2022). Complexes with unavailable TM-helix coordinates or incomplete geometric data were retained as unevaluable rather than excluded.

Inter-protein contact composition was calculated using the same residue-contact definitions as for highTM–highTM complexes. However, for lowTM complexes, the dominant interface type was assigned according to the largest fraction among TM–TM, TM–SOL, and SOL–SOL contacts, with exact ties treated separately. Unlike the highTM–highTM classification, this approach did not require a minimum 50% contribution from the dominant contact category, better accommodating the limited TM interaction surfaces of lowTM proteins and their potentially substantial extramembrane interfaces.

### Membrane penetration of soluble interaction partners

HighTM–soluble heterodimers were identified as complexes containing one protein with at least three annotated TM helices and one partner without annotated TM segments. The membrane boundaries and normal were determined from the highTM protomer, defining its hydrophobic membrane slab. Soluble-partner residues were evaluated according to their positions relative to this slab. For each complex, the percentage of soluble-partner residues located within the hydrophobic core and within the 8 Å membrane-interface region was calculated. Maximum penetration depth was determined from the distance of the deepest-penetrating soluble-partner residue from the nearest hydrophobic-core boundary, measured along the membrane normal. Complexes without core penetration were assigned a depth of zero. Of 4,643 candidate heterodimers, sequence-based chain assignments were obtained for 4,642, and 4,618 complexes with highTM-derived membrane thicknesses of 15–35 Å were retained for quantitative analysis.

### Local AlphaFold runs

AlphaFold-Multimer predictions were performed locally using AlphaFold v2.2.0 with the AlphaFold-Multimer v2 model parameters (Evans et al. 2021). Ten predictions were generated using five models with two predictions per model. AF3 was retrieved from https://github.com/google-deepmind/alphafold3 and used with default parameters (Abramson et al. 2024). Predicted complexes were compared with the experimentally determined rabbit SERCA1a–PLN complex (PDB ID: 4y3u; (Akin et al. 2013)).

The experimentally determined cholesterol-mediated homodimer of the norepinephrine transporter (NET; PDB: 9ke3; (H. Zhang et al. 2025)) was used to investigate the ability of AF3 to reproduce lipid-mediated protein assemblies. Predictions were performed locally using two NET, 14 cholesterol, and two POPG (1-palmitoyl-2-oleoyl-sn-glycero-3-phosphoglycerol) molecules. POPG was used instead of A1EFM ([(2∼{R})-1-[[(2∼{S})-2,3-bis(oxidanyl)propoxy]-oxidanyl-phosphoryl]oxy-3-nonanoyloxy-propan-2-yl] decanoate) present in the experimental structure because its longer tails represent a more physiological setup. An initial set of five AF3 models was generated using standard inference parameters. Sampling was subsequently extended by 100 additional models using 20 random seeds (101–120), with five samples generated per seed. Predicted complexes were compared with the experimental structure after superimposing one protomer and examining the relative placement of the second protomer and the cholesterol-mediated interface.

### Prediction and evaluation of protein–cholesterol complexes using AlphaFold3

Experimentally determined transmembrane protein structures containing cholesterol (CCD identifier: CLR) were identified using the Protein Data Bank (wwPDB consortium 2019). Membrane-associated protein chains were selected using available TM annotations, and their sequences were used to prepare AF3 prediction inputs. Experimental reference structures were prepared by extracting the selected protein chains and associated cholesterol residues from the original structures. Targets were grouped according to their structural release dates relative to the AF3 training cutoff (30 September 2021) and the availability of previously determined homologous structures. Homologs were identified using BLAST with an E-value threshold of 10^-10^, minimum query coverage of 70%, alignment length of at least 120 residues, and sequence identity of at least 30%. Homologous structures were required to have been released before the training cutoff. Post-cutoff targets were divided into three categories: those with a homologous structure containing cholesterol, those with a homologous structure lacking cholesterol, and those without a qualifying homolog.

AF3 predictions were performed using protein sequences and varying numbers of cholesterol molecules represented by the CLR chemical component. Multiple predictions were generated for each target using different random seeds and cholesterol copy numbers, with five structural samples generated per prediction job. Our dataset with 181 monomer target identifiers were randomly selected but provide comparable number of proteins in each category, including 43 before-cutoff, 47 homolog-only, 43 cholesterol-containing-homolog, and 48 no-homolog targets.

Cholesterol-binding pose accuracy was evaluated using the Local Distance Difference Test for Protein–Ligand Interactions (LDDT-PLI), calculated with OpenStructure using a 6 Å interaction neighborhood (Biasini et al. 2013). For each predicted cholesterol molecule, the highest valid LDDT-PLI score among the available experimental cholesterol ligands was selected. Target-level scores were calculated by averaging the available numerical ligand scores across the saved prediction samples, cholesterol-copy-number configurations, and experimental-reference comparisons. Group-level distributions and means were then calculated from the target-level averages, giving each evaluable target equal weight. Missing scores were excluded rather than treated as zero. Of the 181 targets, 173 yielded evaluable LDDT-PLI scores, comprising 43, 46, 40, and 44 targets in the four groups, respectively.

### Scripting, plotting, and molecular visualization

Data retrieval, processing, structural analysis, and visualization were performed using Python scripts. Computational workflows were organized using Snakemake to manage dependencies, execution, and reproducibility. Figures were generated using Matplotlib, and molecular structures were visualized using PyMOL. OpenAI Codex was used to assist script development and review, while ChatGPT was additionally used to support the evaluation of selected analytical procedures and outputs. The computational workflows, source code, and associated parameters will be made available at GitHub, with analysis datasets deposited at Zenodo.

