## Supplemental Figures and Tables for "Membrane Geometry Reveals Strengths and Limitations of AlphaFold-Predicted Transmembrane Protein Complexes"

### SUPPLEMENTARY MATERIAL

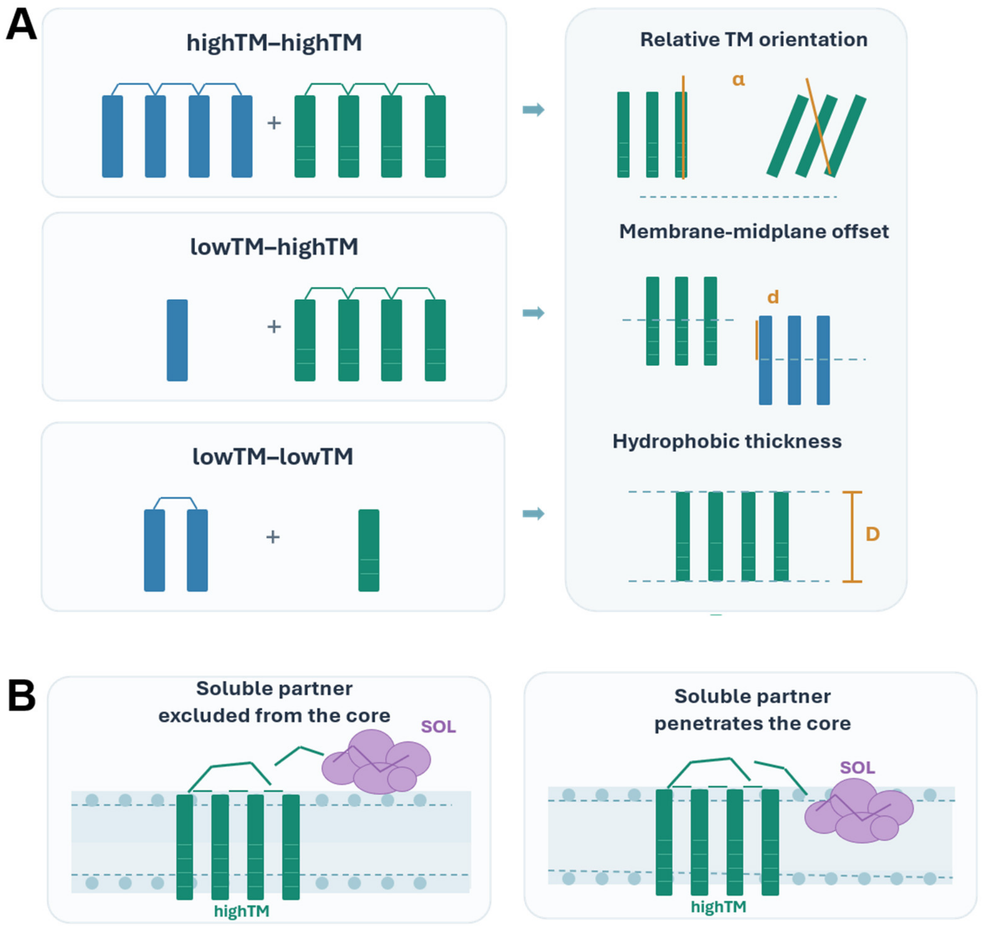


**Supplementary Figure S1. Schematic overview of the geometric analysis of TM–TM (A) and TM–soluble (B) protein dimers.** highTM: proteins containing three or more transmembrane helices; lowTM: proteins containing one or two TM helices; d: hydrophobic thickness; α, relative angle between the membrane orientations of the two protomers.

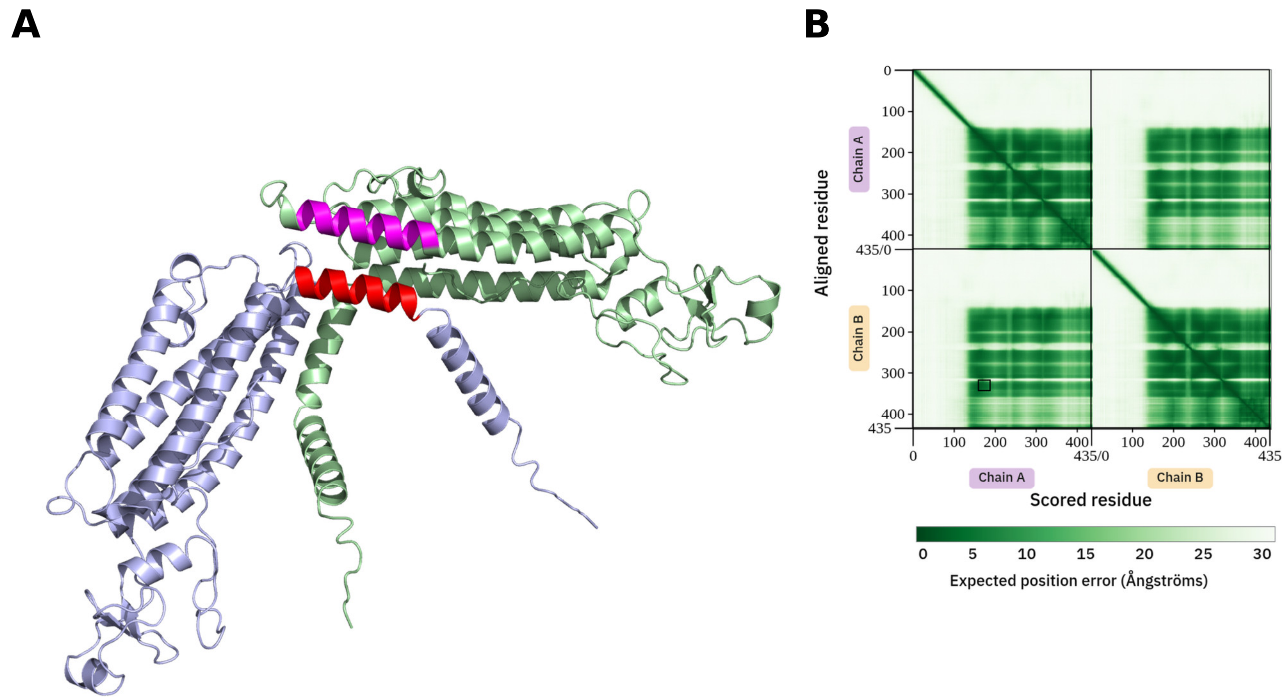


**Supplementary Figure S2. The suspicious intermolecular interaction in the E3 ubiquitin-protein ligase RNFT1 homodimer is associated with low PAE values.** RNFT1 shows an unusual predicted transmembrane topology, and the two protomers adopt a suspicious, approximately perpendicular arrangement rather than the expected parallel membrane orientation (Figure 2). **(A)** The local confidence of the intermolecular interaction was assessed using predicted aligned error (PAE). The interacting regions comprise residues 155–167 in chain A (red) and residues 320–333 in chain B (magenta). **(B)** The corresponding interchain PAE values for these residue pairs are low, indicating that AlphaFold3 predicts their relative positioning with high confidence. The relevant region of the PAE matrix is marked by a black rectangle.

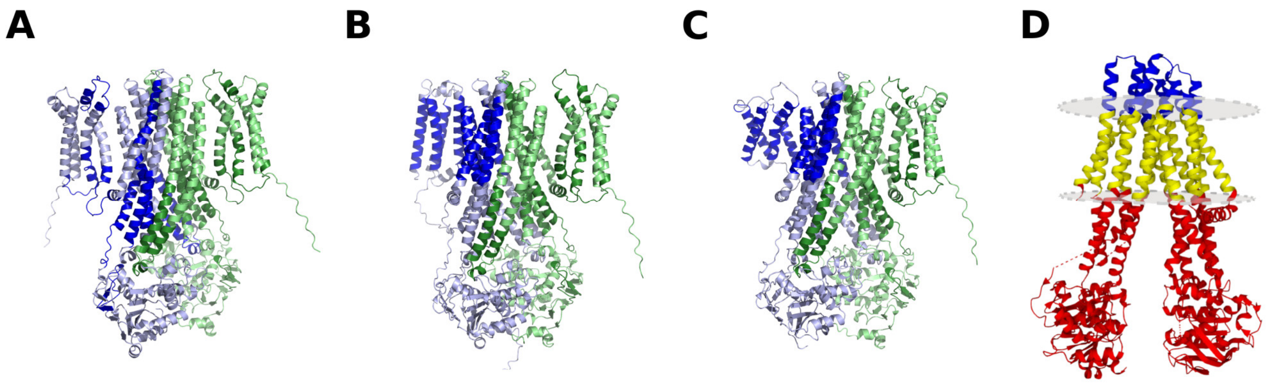


**Supplementary Figure S3. Two of the three AFDB TAP1-containing complexes are unlikely to represent physiological assemblies. (A)** The predicted TAP1–TAP1 homodimer is structurally well resolved, but shows a large membrane-midplane displacement and is therefore classified as highly suspicious, primarily because of incorrect TM-region prediction (dark blue and dark green). More importantly, TAP1 homodimerization is not supported by experimental evidence [PMID: 12047747], and a TAP1–TAP1 complex would not reproduce the asymmetric ATPase-site architecture required for functional TAP transport [PMID: 17018292]. **(B)** The TAP1–ABCB9 complex is likewise classified as highly suspicious because of mispredicted TAP1 TM regions. Although TAP1 and ABCB9 are homologous ABC half-transporters, their heterodimerization has not been experimentally established [PMID: 15518576]. Moreover, TAP1 functions predominantly in the ER membrane, whereas ABCB9/TAPL is primarily associated with lysosomal membranes [PMID: 16525794]. **(C)** The physiological TAP1–TAP2 heterodimer is also classified as suspicious by the membrane-geometry analysis because of inaccurate TM-region prediction, despite the overall complex architecture being biologically correct. **(D)** AF3 predicts an inward-closed conformation with closely associated nucleotide-binding domains (NBDs), whereas cryo-EM structures have captured the TAP1–TAP2 heterodimer in an inward-open conformation. One such experimental structure (PDBID: 8t4h) is shown with intracellular (red), extracellular (blue), and transmembrane (yellow) regions assigned by the structure-based membrane-topology method TMDET.

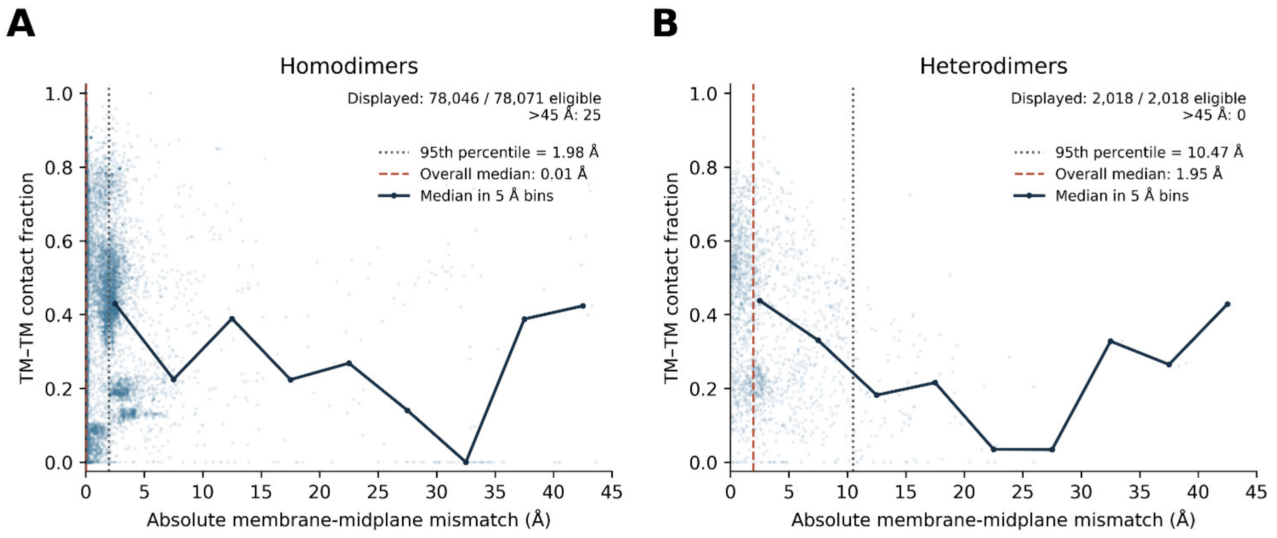


**Supplementary Figure S4. Membrane-midplane mismatch in highTM–highTM homodimers and heterodimers. (A)** Homodimers and **(B)** heterodimers were analysed separately to compare the agreement between membrane positions inferred independently from the two protomers. Light blue points represent individual complexes, and the dark line indicates the median TM–TM contact fraction in 5 Å mismatch bins. Vertical dashed lines mark the overall median (red) and the 95th percentile (gray) of the mismatch distribution. The heterodimer distribution is substantially broader than the homodimer distribution, with a 95th percentile of 10.47 Å compared with 1.98 Å for homodimers. For visual clarity, complexes with membrane-midplane mismatch >45 Å are omitted from the plot, but were included in percentile calculations.

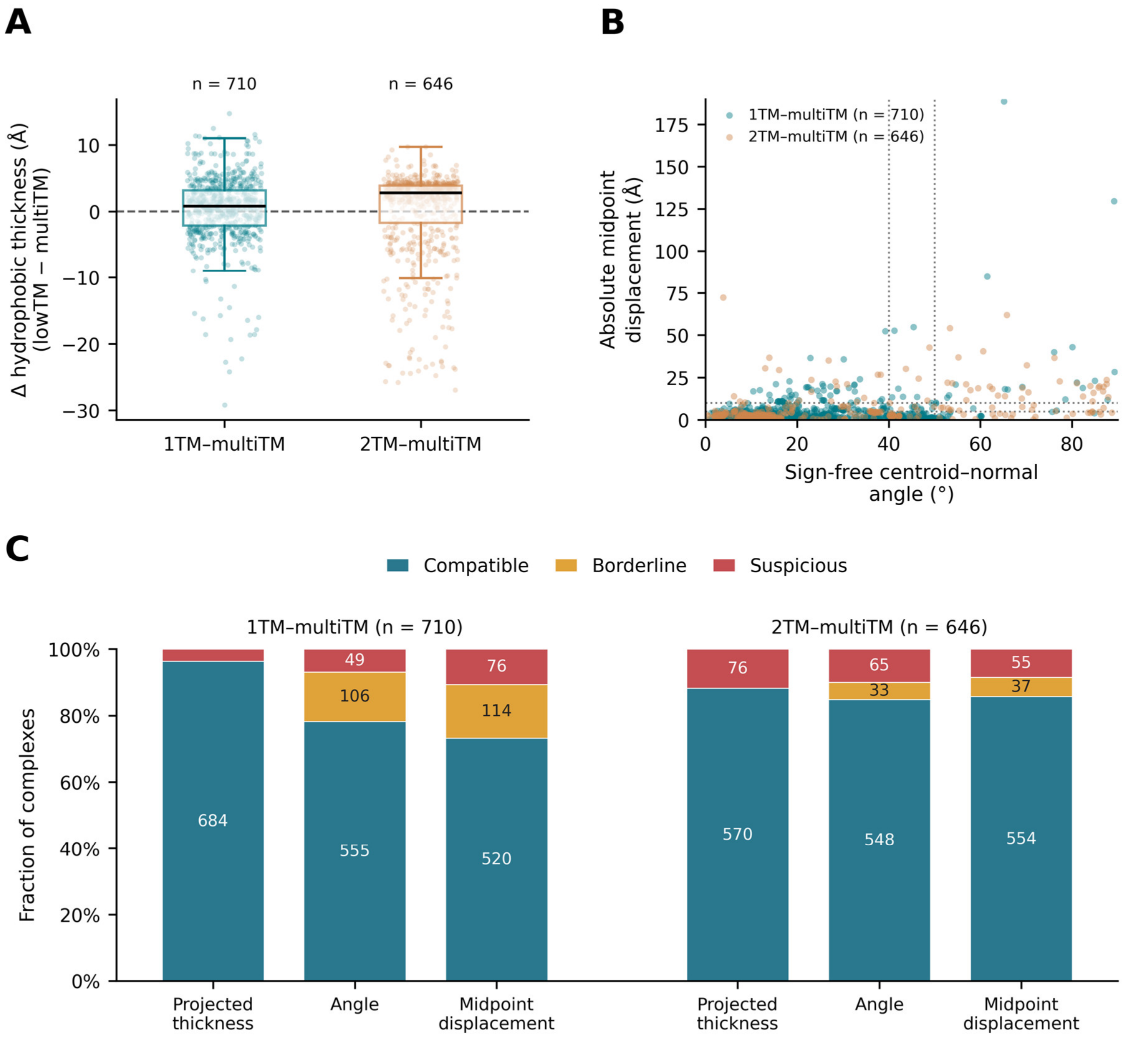


**Supplementary Figure S5. Geometry-based characterization of lowTM–multiTM complexes.** Analyses were performed on the QC-eligible cohort of 1,356 complexes, comprising 710 1TM–multiTM and 646 2TM–multiTM interactions. **(A)** Distribution of signed hydrophobic thickness mismatch, defined as projected lowTM hydrophobic thickness minus reference multiTM hydrophobic thickness. Boxplots are overlaid with jittered individual complexes, and the horizontal zero line indicates perfect agreement between the lowTM projected span and the membrane thickness inferred from the multiTM partner. **(B)** Relationship between the sign-free centroid–normal angle and the absolute midpoint displacement for 1TM–multiTM and 2TM–multiTM complexes. Dashed guide lines indicate the geometry-QC thresholds used for classification: 40° and 50° for angle, and 5 Å and 10 Å for midpoint displacement. **(C)** Criterion-specific QC composition for each topology, shown separately for projected thickness, angle, and midpoint displacement. Bars indicate the fractions classified as Compatible,

Borderline, or Suspicious for each metric. Under the current definition, projected thickness contains only Compatible and Suspicious cases, whereas angle and midpoint displacement include all three categories. Together, these analyses show how 1TM and 2TM lowTM partners differ in membrane-span agreement and in the geometric features that contribute to overall QC assignment.

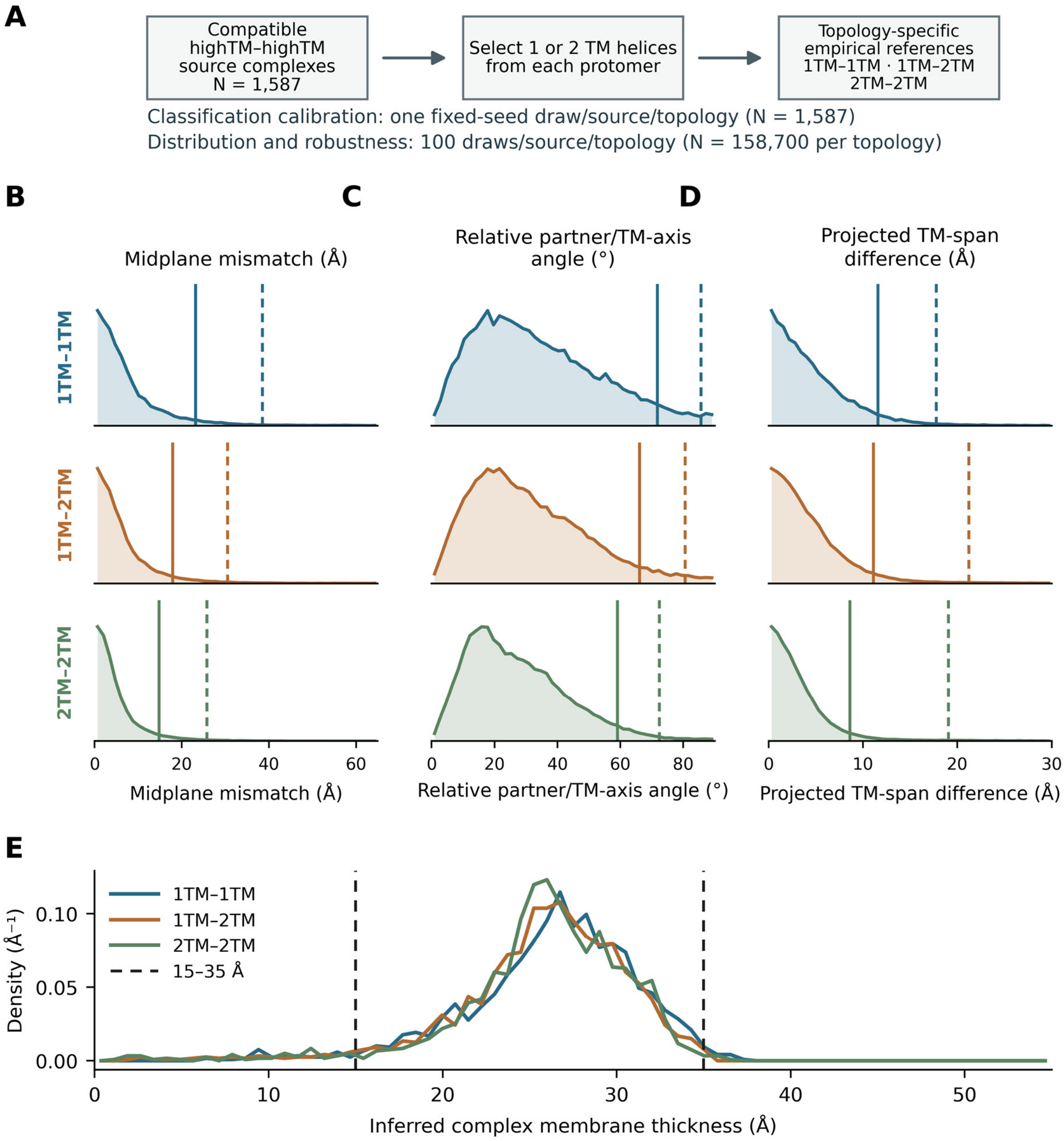


**Supplementary Figure S6. Empirical calibration of lowTM–lowTM membrane geometry. (A)** Schematic of the empirical reference construction. Because one- and two-pass membrane proteins do not contain sufficient TM helices to independently define extracellular and intracellular membrane boundary planes, geometrically compatible highTM–highTM complexes were used as a reference. From 1,587 compatible complexes, one or two TM helices were sampled from each protomer to generate pseudo-1TM–1TM, pseudo-1TM–2TM, and pseudo-2TM–2TM representations. Repeated sampling (100 realizations per source complex and topology; 158,700 observations per topology) was used to characterize the expected distributions and sampling variability. An independent fixed-seed sample containing one pseudo-complex per source complex and topology was used for classification calibration. **(B-D)** Empirical distributions of (B) TM-midplane mismatch, (C) relative partner/TM-axis angle, and (D) projected TM-span difference for the three pseudo-lowTM topologies. Dashed and dash-dotted vertical lines indicate the corresponding topology-specific 95th and 99th percentile reference thresholds derived from the independent calibration sample. **(E)** Distribution of the inferred membrane-spanning thickness, defined as the smaller of the two projected partner TM spans, in the independent calibration sample (N = 1,587 per topology). Vertical dashed lines at 15 and 35 Å indicate the inclusive membrane-thickness interval used as an additional geometry-quality criterion. The dense repeated samples shown in B–D were used to characterize the robustness and expected variation of lowTM geometry and were not treated as independent complexes for classification.

**Supplementary Table S1. Cohort sizes, selection criteria, and corresponding analyses.**

| Dataset or analysis subset | N | Selection criteria or description | Corresponding analysis |
|---|---|---|---|
| **TM–TM complexes: initial cohort** | **190,548** | Both partners annotated as TM proteins with at least one TM helix; 184,521 homodimers and 6,027 heterodimers. Records without usable coordinates retained. | Initial TM–TM dataset |
| TM–TM complexes with evaluable contacts | **189,469** | Valid structural coordinates and inter-protein contact fractions; 183,442 homodimers and 6,027 heterodimers. | Figure 1 |
| **HighTM–highTM complexes with evaluable geometry** | **125,657** | At least three structurally usable TM helices of ≥15 residues per protomer; 122,553 homodimers and 3,104 heterodimers. | Figure 2 |
| HighTM–highTM complexes with assigned interface classes | **125,564** | Evaluable relative TM-axis angle and assigned TM, Mixed, or SOL interface class. | Figure 2D |
| HighTM–highTM complexes with evaluable midplane mismatch | **80,089** | Relative TM-axis angle ≤30°, and valid membrane-midplane mismatch and contact fraction; 78,071 homodimers and 2,018 heterodimers. | Figure 2C; Supplementary Figure S4 |
| HighTM–highTM complexes displayed in mismatch plots | **80,064** | Same evaluable cohort, excluding 25 complexes with mismatch >45 Å for visualization only. | Figure 2C; Supplementary Figure S4 |
| **LowTM–highTM candidate complexes** | **1,591** | One partner with one or two TM helices and the other with at least three TM helices. | Initial Figure 3 dataset |
| LowTM–highTM complexes passing membrane-thickness QC | **1,356** | HighTM reference thickness within 15–35 Å; 710 1TM–highTM and 646 2TM–highTM complexes. | Figure 3 |
| **HighTM–highTM calibration reference complexes** | **1,587** | Selected from a highTM–highTM cohort classified as geometrically compatible (relative angle ≤40° and minimum individual thickness ≥15 Å). | Pseudo-lowTM calibration |
| Pseudo-lowTM repeated samples | **476,100** | 100 random TM-helix selections per reference complex for each of three topology classes. | Supplementary calibration analysis |
| Pseudo-lowTM threshold calibration samples | **4,761** | One fixed-seed sample per source complex and topology (1,587 per topology), used to derive P95/P99 thresholds. | Supplementary calibration analysis |
| **LowTM–lowTM complexes** | **610** | 459 1TM–1TM, 62 1TM–2TM, and 89 2TM–2TM complexes. | Figure 4 |
| LowTM–lowTM complexes with evaluable geometry | **540** | Required TM-helix coordinates and geometric descriptors available; 70 additional complexes remained unevaluable. | Figure 4A,C |
| **HighTM–soluble candidate heterodimers** | **4,643** | One highTM partner and one partner without annotated TM segments. | Initial Figure 5 dataset |

| Dataset or analysis subset | N | Selection criteria or description | Corresponding analysis |
|---|---|---|---|
| HighTM–soluble complexes with chain assignments | **4,642** | Sequence-based chain assignment successfully completed. | Membrane-penetration analysis |
| HighTM–soluble complexes passing thickness QC | **4,618** | HighTM membrane thickness within 15–35 Å. | Figure 5 |
| **NET–cholesterol AF3 predictions** | **105** | Five initial models and 100 additional samples using 20 random seeds. | Figure 7A–C |
| **Cholesterol-binding benchmark targets** | **181** | Selected experimental membrane protein–cholesterol targets, grouped by training cutoff and homolog availability. | Initial Figure 7D dataset |
| Cholesterol-binding targets with evaluable scores | **173** | Valid LDDT-PLI scores; four groups containing 43, 46, 40, and 44 targets. | Figure 7D |

**Abbreviations.** TM, transmembrane; highTM, protein containing at least three TM helices; lowTM, protein containing one or two TM helices; QC, quality control; LDDT-PLI, Local Distance Difference Test for Protein–Ligand Interactions. The initial TM–TM dataset retains unresolved coordinate records; figure-specific subsets reflect additional contact-data, coordinate, or geometric requirements. Repeated pseudo-lowTM samples are not independent protein complexes.
